# Spontaneous breaking of symmetry in overlapping cell instance segmentation using diffusion models

**DOI:** 10.1101/2023.07.07.548066

**Authors:** Julius B. Kirkegaard

**Affiliations:** Department of Computer Science & Niels Bohr Institute, Unviersity of Copenhagen, Copenhagen, Denmark

## Abstract

Instance segmentation is the task of assigning unique identifiers to individual objects in images. Solving this task requires breaking the inherent symmetry that semantically similar objects must result in distinct outputs. Deep learning algorithms bypass this break-of-symmetry by training specialized predictors or by utilizing intermediate label representations. However, many of these approaches break down when faced with overlapping labels that can appear, e.g., in biological cell layers. Here, we discuss the reason for this failure and offer a novel approach for instance segmentation based on diffusion models that breaks this symmetry spontaneously. Our method outputs pixel-level instance segmentations matching the performance of models such as cellpose on the cellpose fluorescent cell dataset while also permitting overlapping labels.

---

Cell instance segmentation is often the first ingredient in the data processing pipeline of computational assays in cell biology. In contrast to semantic segmentation, instance segmentation requires individual labeling of cells, a procedure that is particularly challenging when cells are densely packed. Recent advances in deep learning approaches to cell instance segmentation have demonstrated how to handle cell layers that lie adjacent to one another [1], [2], [3], [4], which is difficult to handle by pioneering approaches such as UNet [5]. However, under some conditions, cells might not only pack closely at high packing densities but even start overlapping. In this brief paper, we discuss the implications of overlaps in data on existing methods and present a novel architecture based on diffusion models that natively handles overlap.

Semantic segmentation is the task of assigning each pixel in an image a semantic value. In the simplest case of cell layers, this amounts to separating the background pixels from cell pixels [Fig. 1A]. Training a semantic segmentation model is simple because the output of the model can be used directly as labels for the training. In contrast, instance segmentation requires breaking the object-object symmetry by assigning each object a unique label [Fig. 1A]. Such random labeling should not be used as training labels, as there is no inherent truth to their values. Instead, instance segmentation models exploit intermediate representations for the labels.

**Fig. 1.**
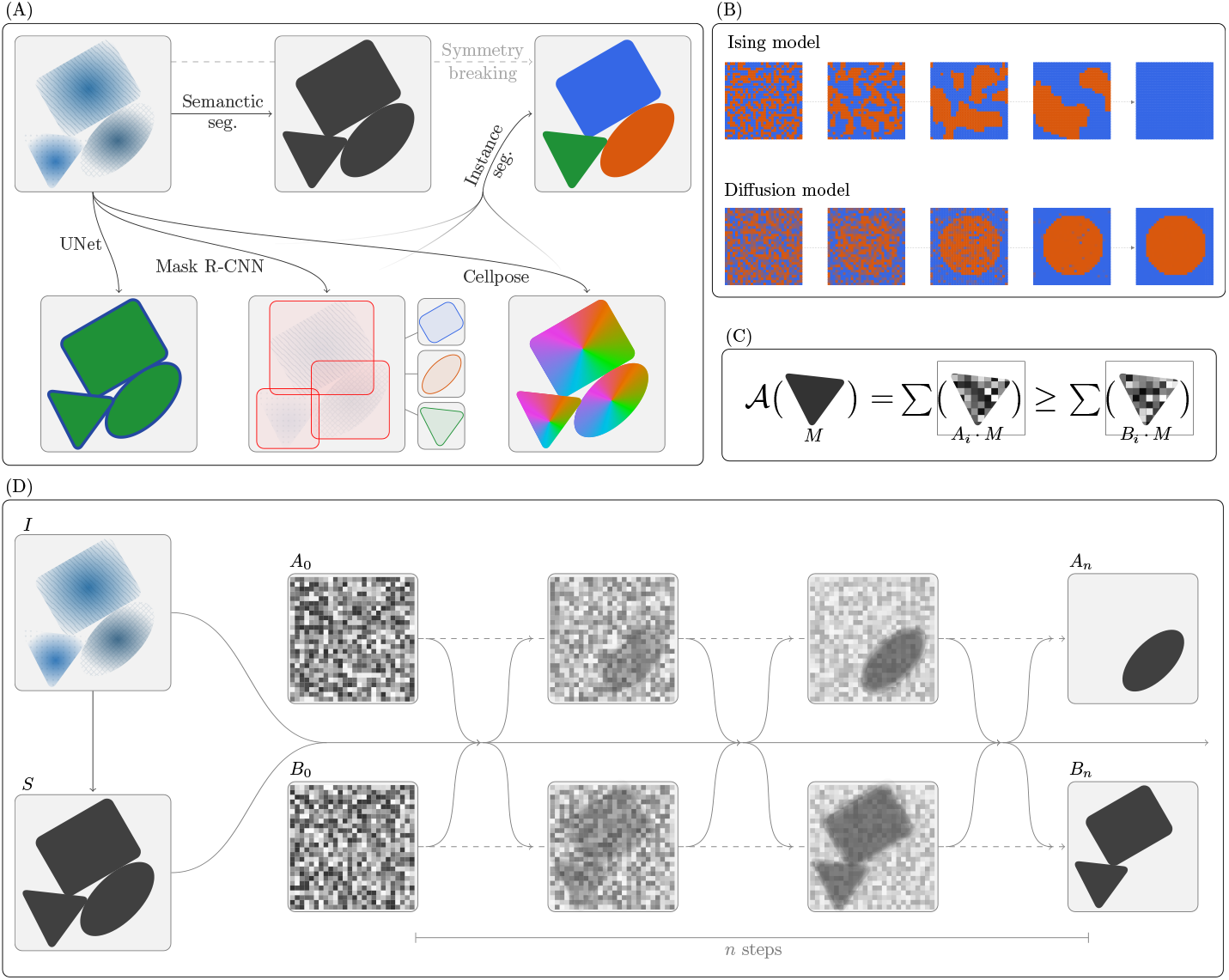
Instance Segmentation. (A) Illustration of an image and its semantic segmentation and instance segmentation, as well as three models for calculating an instance segmentation. UNet predicts a field that ensures small edges between objects making instance segmentation possible by simple flood fill. Mask-R-CNN predicts bounding boxes and associated masks directly at a fixed resolution. Cellpose predicts gradient fields whose basins-of-attraction are instance masks. (B) An Ising model initialized with a noisy field eventually collapses to a single overall mode. A diffusion model is a denoising process starting from noise. (C) In the diffusion split model, a mask *M* belongs in the A-split if the sum of the pixels within the mask is larger in *A*_*i*_ compared to *B*_*i*_. (D) The diffusion split model. An image *I* and its semantic segmentation *S* (e.g. output of UNet model) is the input to the diffusion model along with two noisy frames *A*_0_, *B*_0_. The denoising diffusion process spontaneously assigns instance masks the A- or B-split over *n* steps.

The most direct approach is that used by UNets, which employ semantic segmentation labels that ensure a gap between touching cells [5], as illustrated in Fig. 1A. To achieve this behavior, the models are trained with large weights/attention given to the edges. The obvious downside to this approach is that the correctness of the overall prediction depends on single pixels. A separate approach that has seen wide adoption is Mask-R-CNN [6]. This method predicts YOLO-style [7] bounding boxes for each instance and separately predicts masks for each predicted box [Fig. 1A]. This approach is extremely versatile but has the downside that mask prediction happens at a fixed resolution: the instance segmentation can therefore not become accurate on a pixel level. Furthermore, in contrast to UNet, Mask-R-CNN also breaks the symmetry of the instances, as the output (of each predictor head) is an ordered list of predictions, and this ordering is arbitrary. Thus Mask-R-CNN models must specialize their predictors. The best of both worlds is achieved by the cellpose model [2]. In cellpose, labels are gradient fields of the distance map to the center of cells [Fig. 1A] from which instance segmentations can be achieved by simply calculating basins-of-attraction. This method thus performs pixel-level predictions while avoiding the issue of depending on pixel-thin edges, which in turn leads to significant improvements in accuracy.

What happens when the data include overlapping cells that must be resolved into complete cell masks? As pixel-level predictors, such as UNet and Cellpose, assign individual pixels to instance masks, these methods cannot be used for overlapping data. On the other hand, the Mask-R-CNN approach can make overlapping predictions because the separate mask predictions happen individually per bounding box — at the cost of worse predictions. The question then remains if the pixel-level advantages of cellpose can be kept while allowing for overlap. To achieve pixel-level predictions for overlapped objects it is clear that more than one prediction per pixel is needed. How can this be achieved in a way that maintains the symmetry between cells? Here, we explore the idea of breaking this symmetry *spontaneously*.

Spontaneous breaking of symmetry is a well-known phenomenon in statistical physics, where for instance in the Ising model, initial noise leads to the symmetry breaking of the overall system [Fig. 1B]. Such an approach can be applied to an instance segmentation task if noise spontaneously chooses whether or not to include an instance mask by running an Ising-like model in each mask. To make this idea concrete and trainable, we employ diffusion models [8], which are a class of sampling methods that exploit an initial random field of noise and train a Markov Chain denoising process [Fig. 1B]. Diffusion models have previously been shown to be adaptable to the task of semantic segmentation [9]. The input to our model consists of the image to be segmented *I*, a semantic segmentation of the background/foreground *S*, and two random initializations *A*_0_ and *B*_0_ [Fig. 1D]. We then run a Markov Chain denoising process

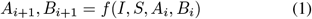

for *n* steps, which spontaneously splits the semantic segmentation *S* into two parts in such a way that instance masks are either fully stored in *A*_*n*_ or *B*_*n*_ [Fig. 1D]. Such a model can then be run recursively, with *A*_*n*_ and *B*_*n*_ going in the place *S* in subsequent runs, to end up with a full instance segmentation [Fig. 2A].

**Fig. 2.**
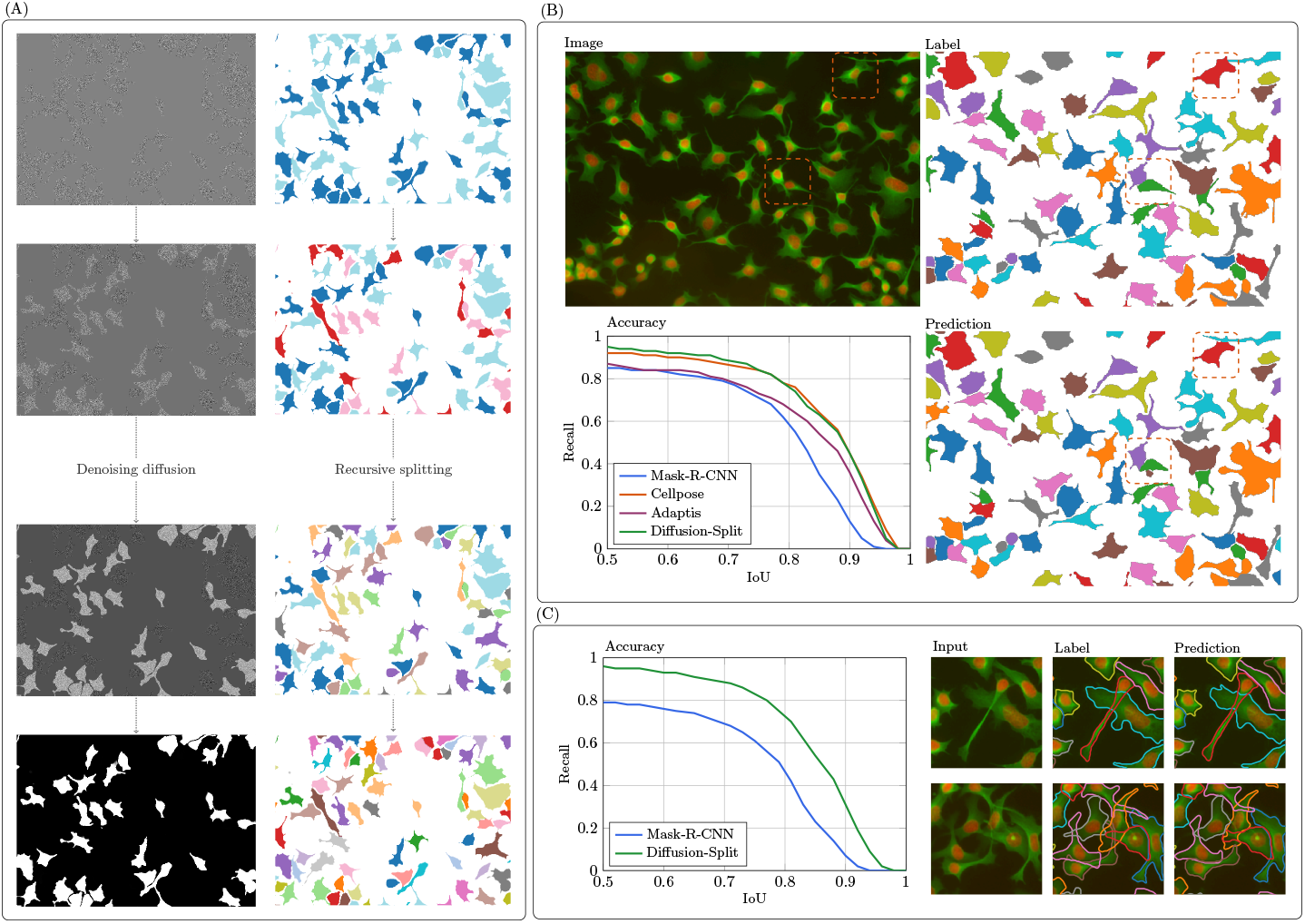
Evaluation of diffusion split. (A) Emergence of the A/B-split from noise (left) and recursive splitting into individual instance masks (right). (B) Example image from the cellpose dataset [2] with two examples of overlap marked, with corresponding labels that ignore this overlap. The plot shows the accuracy of Mask-R-CNN, cellpose, Adaptis, and our method on this dataset, with an example prediction of our method shown as well. Accuracy (in terms of recall) is plotted as a function of an Intersection-over-Union threshold. (C) Results on a relabelled cellpose dataset that accounts for overlap. Only Mask-R-CNN and the present method can output overlaying masks.

The difficulty in the above specification lies in training the model in a way that ensures that masks remain fully in either the *A* or *B* split. Here, we take inspiration from the Ising model’s tendency to collapse to the value chiefly implied by the noise. Thus, during training, a mask *M* is labeled for the *A* split if ∑*A*_*i*_ *· M* ≤ ∑*B*_*i*_ *· M*, i.e. if the pixel sum over a mask is higher in *A*_*i*_ than in *B*_*i*_, as illustrated in Fig. 1C. Training then follows standard procedures for diffusion models, i.e. sampling of noise (*A*_0_, *B*_0_) and a time 0 *i n*, and then gradient descent with a single cross-entropy loss term. We employ UNets for the neural network architecture, similar to that used by cellpose. The neural network (NN) is tasked with predicting *A*_*n*_, *B*_*n*_ at all stages of the process, partially updating *A*_*i*_, *B*_*i*_ with its prediction by e.g.

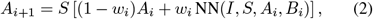

where we choose *w*_*i*_ = (*t*_*i*+1_ − *t*_*i*_)*/*(1 − *t*_*i*_) in which *t*_*i*_ is a predefined log-scaled schedule. In our experiments, we have used 100 steps in the diffusion process with *t*_*i*_ = 10^*i/*10−10^.

To evaluate our approach, we consider the florescent cell data of the cellpose dataset [2]. The images in this dataset do contain overlapping cells [Fig. 2B/C], but the labels ignore this (thus forcing incorrect predictions). We begin by comparing our model to existing models in this setting and subsequently consider a relabelled cellpose dataset that correctly labels overlap.

Model performances are shown in Fig. 2B. In accordance with previous comparisons [2], cellpose demonstrates a superior accuracy at all intersection-over-union thresholds compared with Mask-R-CNN. Our diffusion model split approach achieves approximately the same score as cellpose, thus demonstrating the same improvement over Mask-R-CNN, but with a model that generalizes to overlapping cells.

We then selected the cell images from the cellpose dataset that had the highest degree of overlap and had these relabelled with labels that include all overlap [Fig. 2C]. This is a significantly smaller dataset (2,166 labels, compared to 14,594 in the full), and we thus evaluate our model on this dataset using a 10-fold cross-validation scheme. Retraining Mask-R-CNN on this more complex dataset — both in terms of the more complex predictions needed and less training data — results in a significant decrease in accuracy compared to the standard cellpose dataset [Fig. 2C]. We are unable to run the cellpose model itself on this dataset, as there is no way to have it train on or predict overlapping masks. Running our diffusion model on this data set shows an accuracy that almost maintains the high levels of the original dataset despite the introduction of overlap [Fig. 2C]. Inspecting the preditions, the overlap is correctly resolved in simple overlap [Fig. 2C, top row], but the model struggles in complex settings [bottom row], likely due to the limited amount of training data.

Our approach of using a diffusion model for instance segmentation can be seen as conditioning the predictions on random noise. It is this conditioning that allows pixel-level predictions for overlapping labels. A separate approach is to condition on location, which has previously been studied for instance segmentation [10], [11], albeit not for overlapping labels. For example, Adaptis [10] trains two networks: one network proposes locations and one predicts masks conditioned on those locations. Fig. 2B shows the performance of Adaptis on the cellpose dataset. While it does outperform Mask-R-CNN (at high IoU’s), its accuracy is significantly below that of cellpose and our diffusion approach. While the approach of Adaptis in-principal can predict overlapping masks, it is implemented for panoptic segmentation, which assigns single pixels only one class.

In terms of accuracy, using our diffusion model approach is a plug-in replacement for cellpose that opens the possibility for overlap predictions. Training times are slower than cellpose, however, and the main drawback is in terms of inference times: each prediction requires a number of diffusion denoising processes to be run, in which each step is a full UNet pass, while cellpose only requires a single UNet pass. To speed up our approach, we combine it with standard connected components segmentation which ensures that we only run the model on locations where objects touch or overlap is present. In its current implementation, we run the recursive splitting process a fixed number of times — adaptive stopping could lead to further significant speedup. Yet, regardless of optimizations, this approach will never be able to compete with cellpose in terms of speed.

We have presented a simple approach to adapting diffusion models to solve the problem of overlapping instance segmentation. Our approach shows the strength of maintaining the symmetry between labels as it prevents the need for specialized predictors. Code is available at https://github.com/kirkegaardlab/diffusionsplit, and the overlapping labels for the cellpose dataset at https://github.com/kirkegaardlab/cellpose-overlap.

## Acknowledgments

This project has received funding from the Novo Nordisk Foundation Grant Agreement NNF20OC0062047.

